# A nuclear pore complex component HOS1 mediates UV-B light-induced UVR8 nuclear localization for activating HY5 cascade in *Arabidopsis*

**DOI:** 10.1101/2025.09.22.677739

**Authors:** Shin-Hee Han, Myung Geun Ji, Woe-Yeon Kim, Chung-Mo Park, Jae-Hoon Jung

## Abstract

Upon ultraviolet-B (UV-B) exposure, the photoreceptor UVR8 translocates to the nucleus and interacts with COP1 to activate downstream transcription factors, notably including HY5. However, the mechanism of UVR8 nuclear import has remained unclear^1,2^. Here, we identified HOS1, a component of the nuclear pore complex (NPC), as a key mediator of UVR8 nuclear translocation under UV-B conditions. HOS1 directly interacts with UVR8 specifically in response to UV-B, facilitating its efficient import into the nucleus. Consistent with this role, *HOS1*-deficient mutants exhibit reduced HY5 protein accumulation, compromised UV-B tolerance, and impaired expression of HY5- regulated genes, including those involved in anthocyanin synthesis. These findings reveal a novel role of HOS1 in bridging UV-B perception in the cytoplasm and gene transcriptional activation in the nucleus by enabling UVR8 nuclear import. They highlight a distinct broader role of NPC components in coordinating plant transcriptional response to environmental stimuli.

---

Plants perceive signals from various wavelengths of sunlight, which in turn influence their growth and development. Specifically, ultravioUV-B light, with wavelengths ranging from 280 to 315 nm, elicits various morphological and biochemical changes in plants to confer resistance against UV-B stress^3^. Low levels of UV-B inhibit hypocotyl elongation and induce the synthesis of secondary metabolites, while high levels cause leaf curling, growth inhibition, and ultimately cell death. With the depletion of the Earth’s ozone layer since the Industrial Revolution, the level of UV-B radiation has increased, drawing global attention to research on plant UV-B resistance^4–6^.

Previous studies have identified the photoreceptor UV RESISTANCE LOCUS 8 (UVR8) as the primary sensor through which plants perceive UV-B radiation. Upon activation by UV-B, UVR8 translocates into the nucleus and initiates a cascade of downstream signaling events that lead to plant UV-B responses. A key event in this signaling pathway involves the direct interaction between nuclear-localized UVR8 and the E3 ubiquitin ligase CONSTITUTIVE PHOTOMORPHOGENIC 1 (COP1), resulting in the inhibition of COP1 activity. This interaction prevents the COP1-mediated ubiquitination and degradation of the transcription factor ELONGATED HYPOCOTYL 5 (HY5). Stabilized HY5 subsequently activates the expression of UV-B-responsive genes, including those involved in antioxidant biosynthesis to mitigate oxidative stress^7–10^. Although UVR8 nuclear translocation is essential for activating these protective responses, the molecular mechanisms underlying its nuclear import still remain largely unknown. This study aims to elucidate how HIGH EXPRESSION OF OSMOTICALLY RESPONSIVE GENES 1 (HOS1), a component of the NPC outer-ring complex, mediates the nuclear import of UVR8 under UV-B conditions, thereby triggering the UV-B response through the COP1–HY5 regulatory module.

To investigate the role of HOS1 in UV-B responses, we examined UV-B tolerance in *hos1* loss-of-function mutants. Both *hos1-3* and *hos1-5* mutants exhibited significantly reduced UV-B tolerance compared to wild-type Col-0 plants (Fig. 1a). This phenotype was efficiently rescued by expressing the *HOS1* gene driven by its native promoter in the *hos1-3* mutant background (Supplementary Fig. 1a). HOS1 contains a RING-finger domain essential for its E3 ubiquitin ligase activity, and previous studies have shown that H75Y and C89S substitutions disrupt its enzymatic function^11,12^. Expression of this catalytically inactive HOS1 variant in the *hos1-3* mutant was sufficient to restore UV-B tolerance (Supplementary Fig. 1b), indicating that the ubiquitin ligase activity of HOS1 is not required for its role in UV-B tolerance. These observations suggest that HOS1 contributes to UV-B stress responses through a mechanism independent of its E3 ligase activity.

**Fig. 1:**
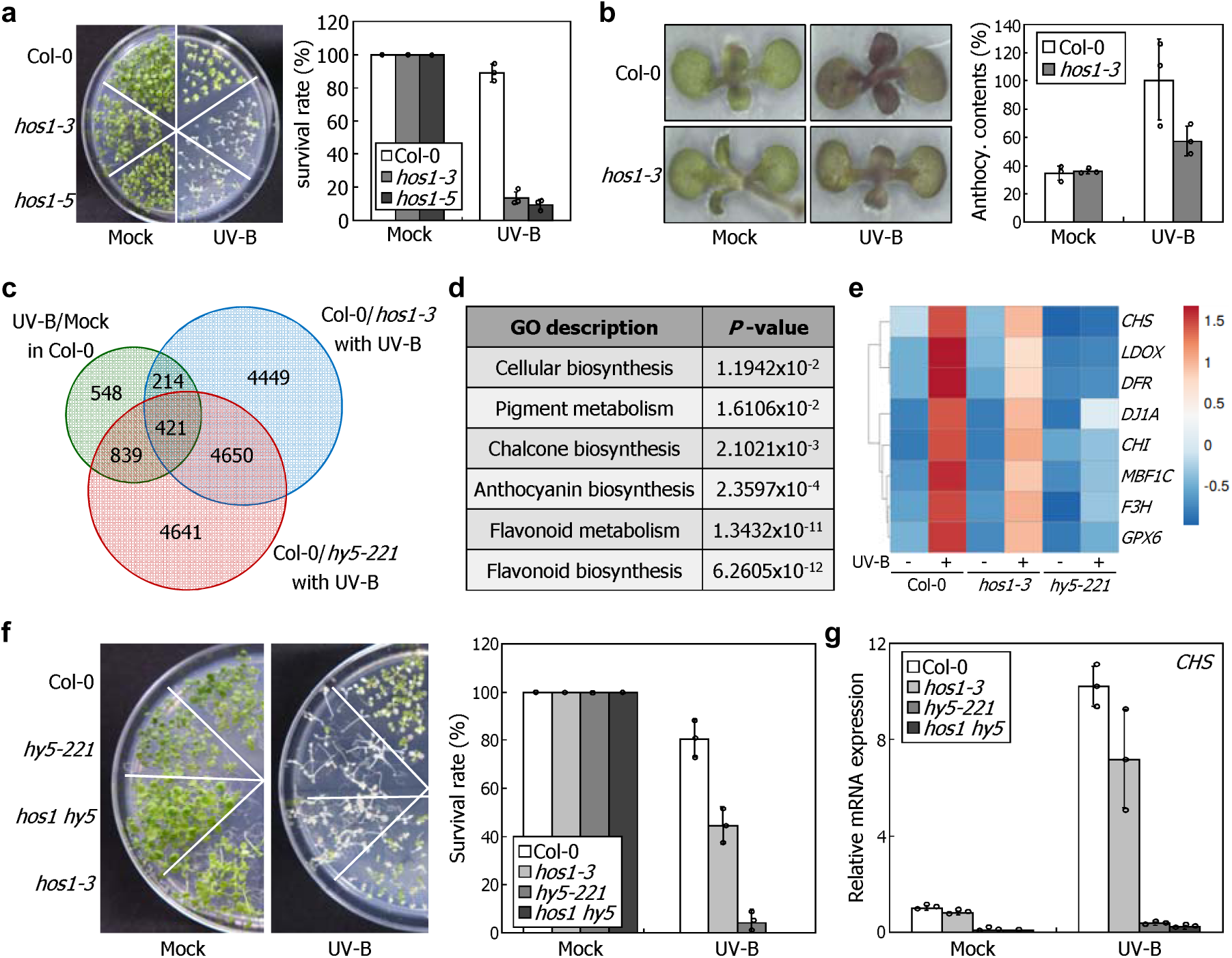
Phenotype of *hos1* mutants hypersensitive to UV-B radiation. Seven-day-old seedlings grown on MS-agar plates at 22℃ were exposed to UV-B radiation for five hours. Three independent measurements, each consisting of 15-20 seedlings, were statistically analyzed. Error bars indicate the standard error of the mean (SE). **a,** UV-B tolerance phenotypes of *hos1* mutants. UV-B-treated seedlings were allowed to recover at 22℃ for 5 d under constant light conditions before measuring their survival rate. **b,** Anthocyanin accumulation in the *hos1-3* mutant. Anthocyanin levels were determined based on absorbance at 537nm, 647nm, and 663nm using cyanidin-3-glucoside as the standard. **c,** Venn diagram showing the overlap of UV-B responsive genes regulated by HY5 and HOS1. Total RNA was extracted from whole seedlings and subjected to RNA sequencing. Differentially expressed genes were selected by the following criteria: genes that exhibited more than a four-fold increase in mRNA expression in Col-0 plants under UV-B conditions and UV-B responsive genes that showed higher expression in Col-0 compared to *hos1-3* and *hy5-221* mutants, respectively. **d,** Gene Ontology (GO) analysis of the 421 overlapping genes in Venn diagram. **e,** Expression profiles of anthocyanin biosynthesis-related genes among the 421 overlapping genes. **f,** UV-B tolerance phenotypes of *hos1-3 hy5-221* double mutant. UV-B treatments and measurements of survival rate were performed, as described in **a**. **g,** Relative mRNA expression levels of *CHS* gene in *hos1-3 hy5-221* double mutant under UV-B conditions.

Plants employ UV-B-induced ROS scavenging mechanisms, such as anthocyanin biosynthesis, to mitigate cellular damages and prevent cell death^6,13^. Our investigation revealed that *hos1-3* mutants accumulated less anthocyanins than Col-0 plants upon UV-B exposure (Fig. 1b), suggesting that HOS1 activates anthocyanin biosynthesis to make plants resilient to oxidative stress under UV-B conditions. To understand how HOS1 is involved in anthocyanin biosynthesis during the UV-B stress responses, we performed RNA sequencing analysis on HOS1-regulated gene expression in response to UV-B. We identified 421 genes that were significantly upregulated by UV-B radiation in Col-0 but not in *hos1-3* or *hy5-221* mutants (Fig. 1c). Gene ontology (GO) analysis of the 421 overlapping genes revealed enrichment in terms related to anthocyanin biosynthesis and metabolic processes, previously shown to be regulated by HY5 under UV-B conditions (Fig. 1d,e, and Supplementary Fig. 2)^14,15^. Notably, anthocyanin-related genes showed similar downregulation patterns in *hos1-3* and *hy5-221* mutants (Fig. 1e and Supplementary Fig. 3), supporting a functional link between HOS1 and HY5.

To examine the genetic relationship between HOS1 and HY5, we generated *hos1-3 hy5-221* double mutant. The double mutant displayed a high level of UV-B sensitivity comparable to that of the *hy5-221* single mutant (Fig. 1f), suggesting that HY5 might function downstream of HOS1 in the same UV-B signaling pathway. Consistently, the UV-B- induced expression of *CHALCONE SYNTHASE* (*CHS*), a well-characterized HY5 target gene, was significantly reduced in the double mutant, similar to the reduction observed in the *hy5-221* mutant (Fig. 1g). These results suggest that HOS1 enhances UV-B tolerance through a HY5-dependent mechanism.

To investigate how HOS1 regulates HY5-downstream gene expression, we first examined *HY5* mRNA and protein levels in *hos1-3* mutants. Under UV-B conditions, *HY5* mRNA levels were comparable between Col-0 plants and *hos1-3* mutants (Fig. 2a). In contrast, UV-B-induced accumulation of HY5 protein observed in Col-0 was abolished in *hos1* mutants (Fig. 2b), showing that HOS1 influence HY5 activity post-transcriptionally by regulating its protein stability rather than transcription under UV-B conditions.

**Fig. 2:**
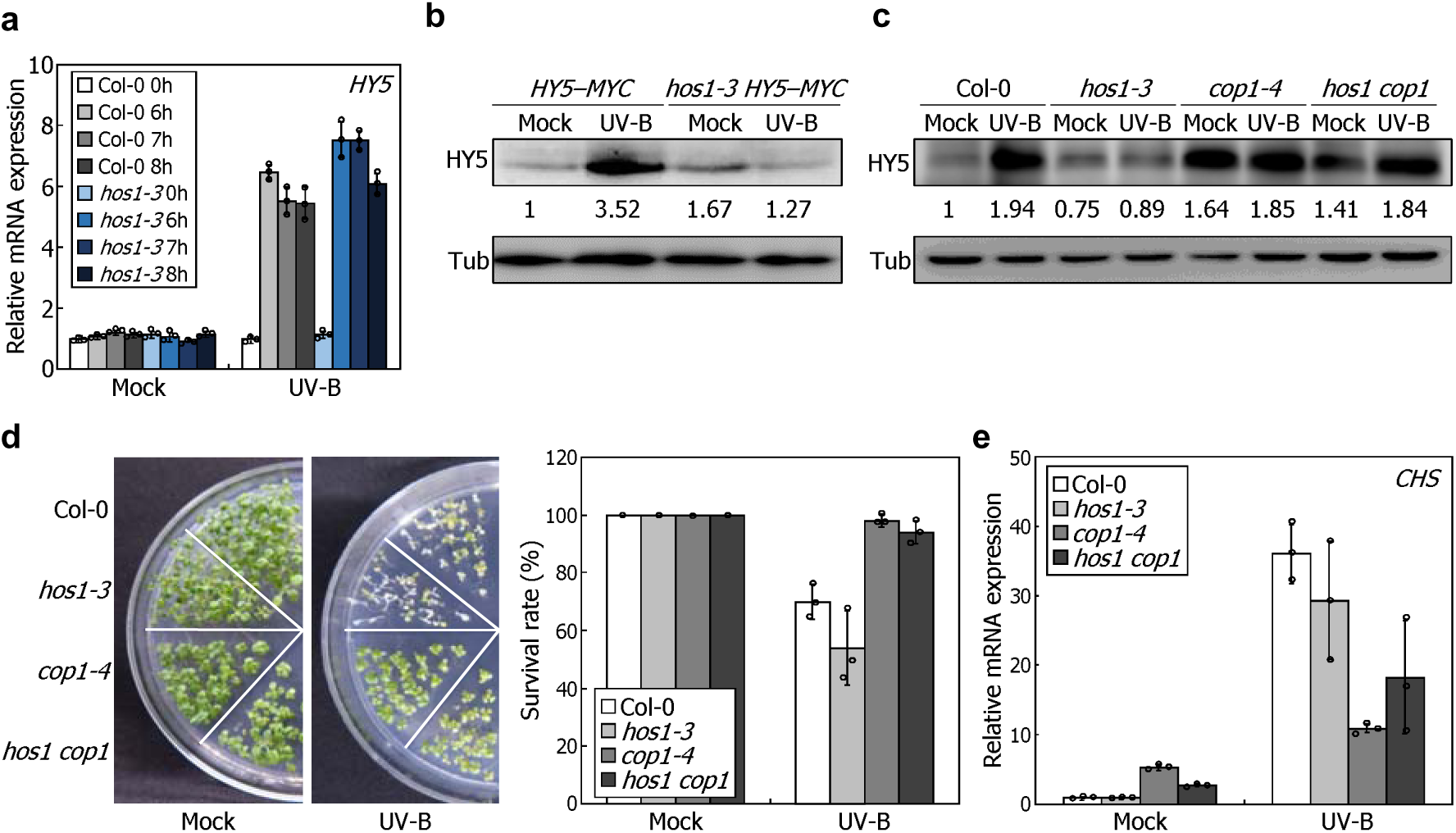
UV-B-induced HY5 accumulation is suppressed in *hos1* mutants. **a,** Relative mRNA expression levels of *HY5* gene in *hos1-3* mutants. Seven-day-old seedlings grown at 22℃ under LDs were exposed to UV-B for the indicated duration before harvesting whole seedlings for total RNA extraction. Levels of mRNA were analyzed by RT-qPCR. Biological triplicates were statistically analyzed for each sample. Error bars indicate the standard error of the mean (SE). **b,** Protein abundance of HY5 in *hos1-3* mutants. *pHY5::HY5–MYC* (*HY5–MYC*) and *hos1-*3 *HY5–MYC* seedlings grown for 7 days at 22℃ under LDs were exposed to UV-B for 5 hours before harvesting whole seedlings for immunoblot analysis. Protein levels of HY5 and tubulin were immunologically detected using anti-MYC and anti-tubulin antibodies, respectively. **c,** HY5 protein accumulation in *hos1-3 cop1-4* double mutant. Immunoblot assays were performed, as described above, except for using anti-HY5 antibody. **d,** UV-B tolerance phenotypes of *hos1-3 cop1-4* double mutants. UV-B treatment and measurements of survival rate were performed, as described in Fig. 1. **e,** RT-qPCR analysis of *CHS* mRNA expression in *hos1-3 cop1-4* double mutants. RT-qPCRs were performed, as described in Fig 1.

Given that COP1 is known to mediate HY5 protein degradation under normal light conditions^16^, we examined whether the failure of HY5 protein accumulation in the *hos1* mutant under UV-B conditions is dependent on COP1 activity. HY protein levels did not increase in the *hos1-3* mutant upon UV-B exposure, whereas in the *hos1-3 cop1-4* double mutant, HY5 protein levels remained elevated, similar to those observed in the *cop1-4* single mutant (Fig. 2c) ^16,17^. Consistently, the double mutant exhibited UV-B tolerance and *CHS* mRNA expression comparable to those of *cop1-4* mutant (Fig. 2d,e), indicating that HOS1 functions upstream of COP1 in regulating HY5 protein stability under UV-B conditions.

Yeast two-hybrid (Y2H) and co-immunoprecipitation (co-IP) assays revealed no physical interaction between HOS1 and either HY5 or COP1 (Supplementary Fig. 4,5). This raised a question of how HOS1 participates in UV-B signaling. On the basis of the observations that COP1 activity and HY5 protein accumulation in the UV-B signaling pathway are UVR8-dependent, we hypothesized that HOS1 might exert its function through interaction with UVR8. To investigate the genetic relationship between HOS1 and UVR8, we generated a *hos1-3 uvr8-6* double mutant. The double mutant exhibited a reduced UV-B tolerance comparable to that of each single mutant (Supplementary Fig. 6), supporting the idea that HOS1 functions in a UVR8-dependent manner in the UV-B signaling pathway.

Next, we examine the possibility of direct interaction between HOS1 and UVR8. Y2H assays revealed that the medial region of HOS1 interacts with UVR8 specifically under UV-B conditions (Supplementary Fig. 7). This interaction was further confirmed *in planta* through co-IP assays using transgenic plants expressing YFP-tagged UVR8, in which HOS1– UVR interaction was detected only after UV-B exposure (Fig. 3a). These results show that HOS1 and UVR8 do not interact under basal conditions, but UV-B activation enables their association.

**Fig. 3:**
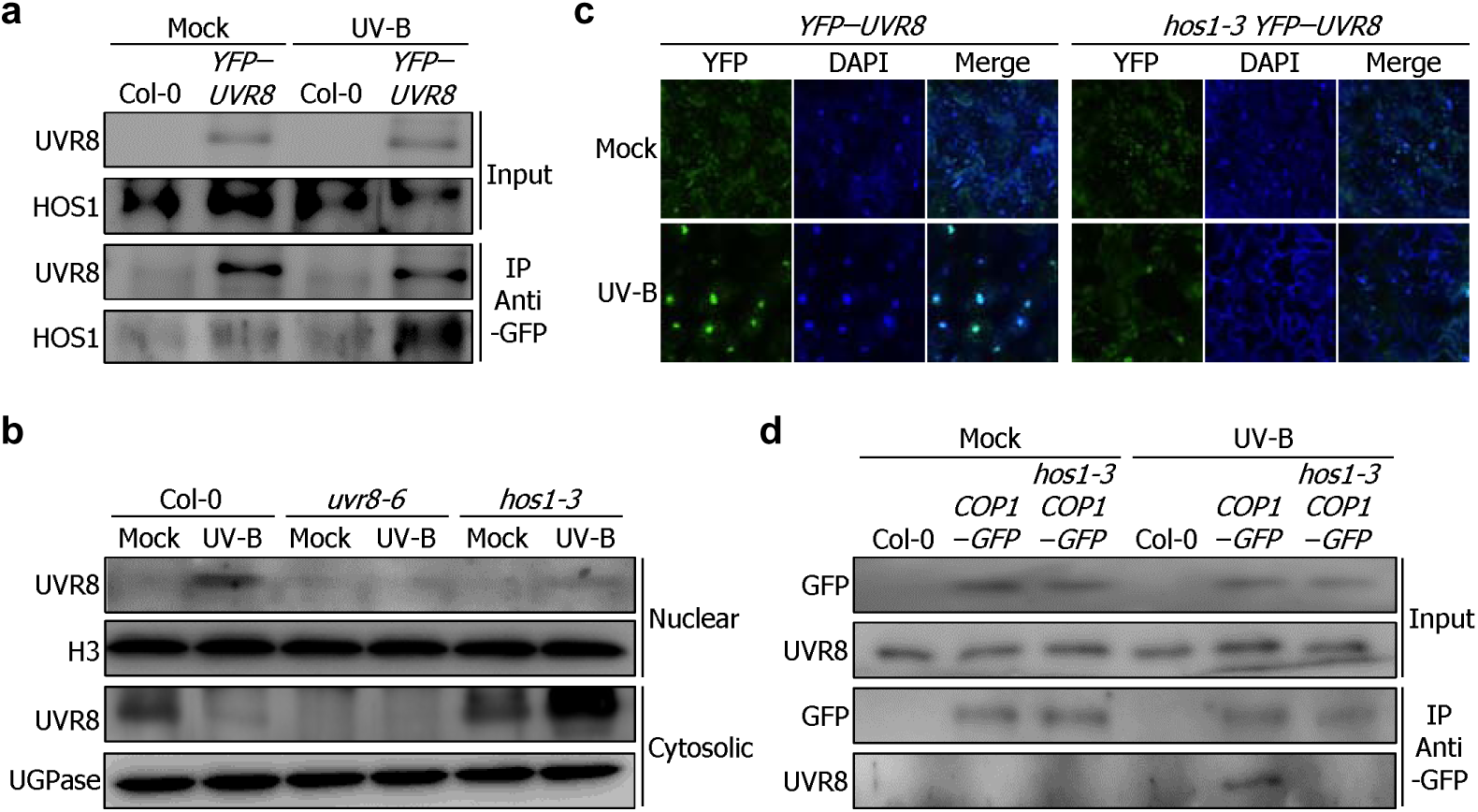
HOS1 mediates the UV-B light-induced nuclear translocation of UVR8. Seven-day-old seedlings grown at 22℃ under LDs were exposed to narrowband UV-B for 1 day before harvesting for assays. **a**, Co-immunoprecipitation (co-IP) analysis of HOS1–UVR8 interaction. Col-0 and *uvr8-6 35S::YFP–UVR8* (*YFP*–*UVR8*) plants were used for co-IP experiments with an anti-GFP antibody. HOS1 and UVR8 proteins were immunologically detected using anti-HOS1 and anti-GFP antibodies, respectively. **b**, Protein abundance of UVR8 in nuclear and cytosolic fractions of *hos1-3* mutants treated with UV-B radiation. UVR8 protein abundance was immunologically detected using an anti-UVR8 antibody. Histone H3 and UGPase served as loading controls for each fraction using anti-histone H3 and anti-UGPase antibodies, respectively. **c**, YFP and DAPI fluorescence changes in *hos1-3* mutants treated with UV-B radiation. *YFP–UVR8* and *hos1-3 YFP–UVR8* seedlings were used for assays. **d**, Co-IP analysis of UVR8–COP1 interaction in the *hos1* mutant. Col-0, 35S::*COP1–GFP* (*COP1–GFP*), and *hos1-3 COP1–GFP* plants were used for co-IP experiments with an anti-GFP antibody. An anti-UVR8 and anti-GFP antibodies were used for the detection of UVR8 and COP1 proteins, respectively.

Recent studies have shown that HOS1, as a component of the NPC outer-ring complex, mediates the nuclear localization of several proteins^18–20^. Given that HOS1–UVR8 interaction occurs specifically under UV-B conditions, we hypothesized that HOS1 facilitates UVR8 nuclear import in response to UV-B. In Col-0 plants, UVR8 resides in the cytoplasm under non-inductive conditions but translocates to the nucleus upon UV-B exposure. Strikingly, in the *hos1-3* mutant, UVR8 fails to accumulate in the nucleus despite UV-B treatment, remaining confined to the cytoplasm (Fig. 3b).

To further confirm the requirement of HOS1 for UVR8 nuclear localization, we employed transgenic plants expressing yellow fluorescent protein (YFP)-tagged UVR8 under the control of the constitutive cauliflower mosaic virus (CaMV) 35S promoter in the *hos1-3* mutant background. As control, we used a complementation line expressing comparable levels of YFP–UVR8 in the *uvr8-6* mutant background^8^. Under UV-B conditions, nuclear-localized YFP–UVR8 fluorescence was clearly observed in control plants, but not in the *hos1-3* mutant background (Fig. 3c). These results demonstrate that HOS1 is required for the UVR8 nuclear accumulation in response to UV-B radiation.

Upon activation by UV-B, UVR8 translocates into the nucleus and interacts with COP1, leading to inhibition of COP1 activity and stabilization of HY5 transcription factor^1,21,22^. Since UVR8 nuclear accumulation is abolished in the *hos1-3* mutant, we investigated whether the UVR8–COP1 interaction also depends on HOS1. In control plants, UVR8 physically interacts with COP1 under UV-B conditions. In contrast, this interaction was not detected in the *hos1-3* mutant background under the same experimental conditions (Fig. 3d). supporting that HOS1 is essential for transmitting the UV-B signals into the nucleus via UVR8.

Recent studies have identified HOS1 as a crucial regulator of plant adaptation to various environmental changes, functioning at the nuclear envelope as a NPC component^23,24^. HOS1 physically interacts with NPC subunits such as NUCLOEPORIN 96 (NUP96) and NUP160, which constitute the NPC outer-ring complex. These two nucleoporins are required for anchoring HOS1 at the NPC, thereby supporting its functional activity in response to environmental stimuli^25,26^.

To determine whether the role of HOS1 in UV-B signaling depends on its localization at the NPC, we analyzed UV-B stress responses in the *nup96-1* and *nup160-3* mutants. Similar to what observed in *hos1* mutants, both *nup96-1* and *nup160-3* mutants exhibited increased sensitivity to UV-B stress (Fig. 4a). Consistently, the expression levels of genes involved in anthocyanin biosynthesis were evidently reduced in the *nup* mutants under UV-B conditions, compared to that observed in Col-0 plants (Fig. 4b). Moreover, HY5 protein levels were significantly decreased in both mutants following UV-B treatment (Fig. 4c). We also found that, unlike in Col-0 plants, UVR8 predominantly accumulated in the cytoplasm of the *nup* mutants even after UV-B exposure (Fig. 4d), a phenotype reminiscent of that observed in the *hos1* mutant. These findings demonstrate that HOS1, via its anchoring at the NPC, is essential for proper UVR8 nuclear import and UV-B signaling transduction in *Arabidopsis*.

**Fig. 4:**
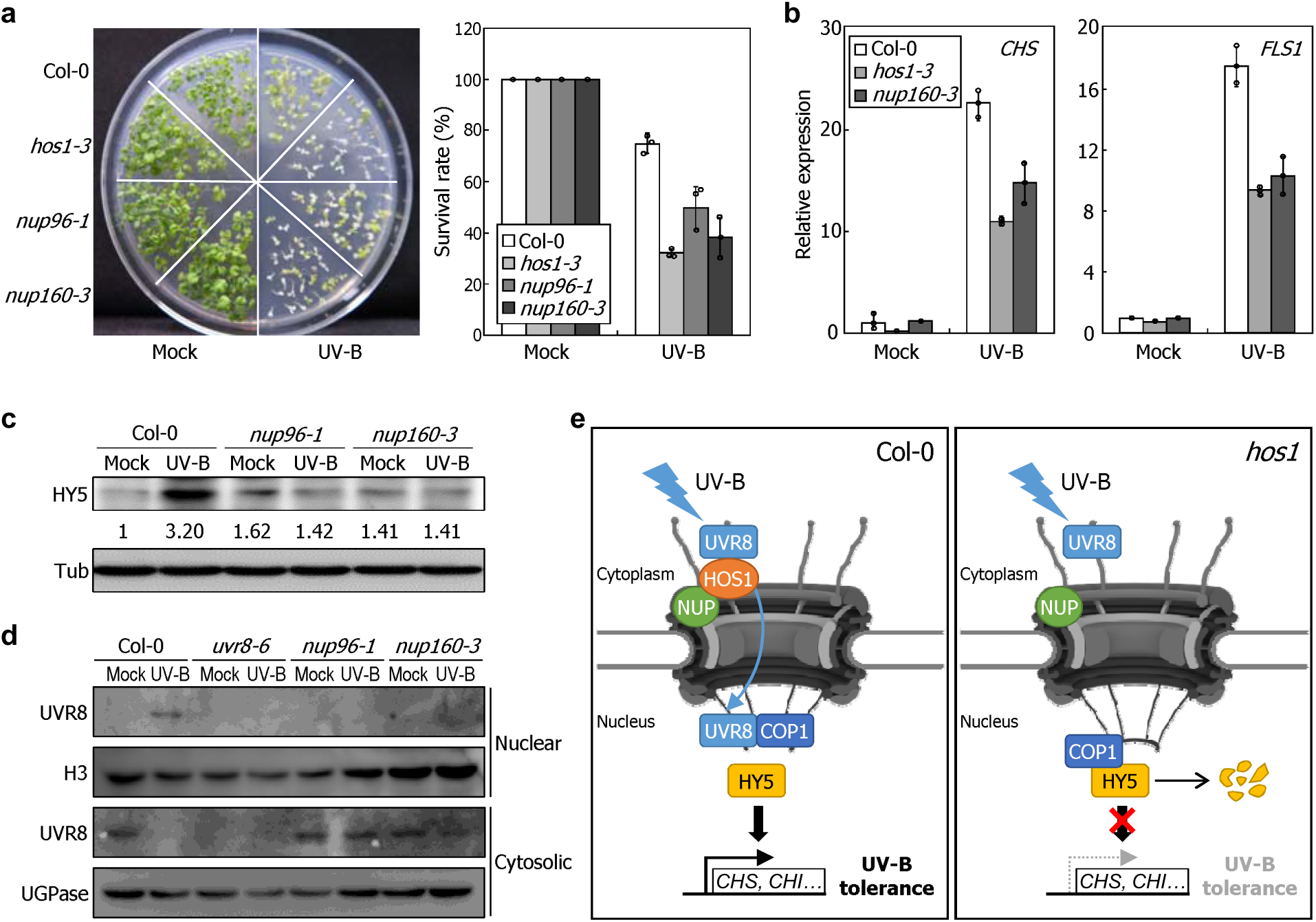
HOS1-interacting nuclear pore complex components NUP96 and NUP160 mediate UV-B light signals for HY5 accumulation. **a**, UV tolerance phenotype of *nup96-1* and *nup160-3* mutants. UV-B treatment and measurements of survival rate were performed, as described in Fig. 1. **b,** RT-qPCR analysis of *CHS* and *FLS1* mRNA expression in *nup96-1* and *nup160-3* mutants. RT-qPCRs were performed, as described in Fig 1. **c**, Protein abundance of HY5 in *nup96-1* and *nup160-3* mutants treated with UV-B radiation. UV-B treatment and immunoblot assays were performed, as described in Fig 3b. **d,** Protein abundance of UVR8 in nuclear and cytosolic fractions of *nup* mutants treated with UV-B radiation. UVR8 protein abundance was immunologically detected using an anti-UVR8 antibody. Histone H3 and UGPase served as loading controls for each fraction using anti-histone H3 and anti-UGPase antibodies, respectively. **e,** Working model for the role of HOS1 in the UV-B signaling pathway. HOS1, as a component of the nuclear pore complex, enhances plant UV-B tolerance by facilitating the nuclear import of UVR8 under UV-B conditions.

Previous studies have reported that HOS1 protein levels increase significantly under heat stress, promoting the expression of DNA repair genes such as RECQ2 to repair heat-induced DNA damage, thereby enhancing plant thermotolerance^27^. In contrast, UV-B exposure did not notably alter the mRNA or protein levels of HOS1, nor did it affect RECQ2 mRNA expression (Supplementary Fig. 8). These results suggest that HOS1 mediates plant responses to UV-B and heat stress through distinct, non-overlapping pathway.

In our working model, HOS1 specifically interacts with UVR8 under UV-B conditions, facilitating its nuclear accumulation. The nuclear-localized UVR8 binds to COP1 and inhibits its activity, leading to stabilization of HY5 and activation of downstream UV-B-responsive genes, including those involved in anthocyanin biosynthesis (Fig. 4e). How does HOS1 mediate UVR8 nuclear import? In the absence of UV-B, UVR8 exists as a cytoplasmic homodimer. Upon UV-B exposure, the dimer dissociates into monomers, which are small enough to pass through nuclear pores^28,29^. While this process has been proposed to occur via passive diffusion, our findings show that UVR8 fails to accumulate in the nucleus in *hos1* mutants, suggesting that NPC subunits including HOS1 play a key role in facilitating UVR8 nuclear import, probably via a facilitated transport system using transport proteins. Future researches could shed light on this aspect.

In conclusion, our study demonstrates that HOS1, functioning as a critical component of the NPC, interacts with UVR8 under UV-B conditions to mediate its nuclear localization. Once translocated into the nucleus, UVR8 activates anthocyanin synthesis via HY5, enhancing UV-B tolerance. Notably, this functioning of HOS1 requires its anchoring at the NPC through interactions with NPC subunits such as NUP96 and NUP160, highlighting a specialized role of the NPC in plant environmental adaptation. These findings propose that the NPC acts not only as a passive gateway but also as a regulatory platform controlling the nuclear access of key signaling proteins in response to environmental cues.

## Acknowledgements

We thank Dr. Roman Ulm (University of Geneva, Switzerland) and Dr. Ruohe Yin (Shanghai Jiao Tong University, China) for providing the *uvr8-6 35S::YFP–UVR8* seeds and Dr. Sang Yeol Lee and Dr. Chang-Ho Kang (Gyeongsang National University, Republic of Korea) for providing the *35S::COP1–GFP* seeds. This work was supported by grants from the National Research Foundation of Korea (NRF) funded by the Korean government (grant nos RS-2025-00573487 and RS-2021-NR060084 to J.-H.J., RS-2022-NR070837 and RS-2025-00555778 to W.-Y.K., and RS-2022-NR075018 to S.-H.H.).

## Author contributions

S.-H.H. and J.-H.J. conceived and designed the experiments. S.-H.H. performed experiments and maintained plant materials. M.G.J. created the anti-HOS1 antibody under the supervision of W.-Y.K. C.-M.P analyzed the data. S.-H.H. and J.-H.J. wrote the paper with the help of W.- Y.K. and C.-M.P. All authors approved the paper for submission.

## Additional information

**Extended data** is available for this paper at XXXX.

**Supplementary information** The online version contains supplementary material available at XXXX

**Correspondence and requests for materials** should be addressed to S.-H.H. or J.H.J.

## Methods

### Plant materials and growth conditions

All *Arabidopsis thaliana* lines used were in Columbia (Col-0) background. Sterilized seeds were cold-imbibed (4□) in darkness for 3 d and allowed to germinate on ½ X Murashige and Skoog-agar (MS-agar) plates under long day conditions (LDs, 16-h light and 8-h dark cycles). White light illumination was provided at light intensity of 120 μmol photons m^-2^ s^-1^ using fluorescent FLR40D/A tubes (Osram). Plants were grown in a controlled culture room set at 22□ with relative humidity of 60%.

The *cop1-4*, *hos1-3* (T-DNA insertion; SALK_069312), *hos1-5* (T-DNA insertion; SAIL_1211_D02), *hy5-221*, *nup96-1* (T-DNA insertion; SALK_109959), *nup160-3* (T-DNA insertion; SALK_126801), and *uvr8-6* (T-DNA insertion; SALK_033468) mutants and *hos1-3 35S::MYC–HOS1*, *hos1-3 35S::MYC–mHOS1*, *35S::YFP–UVR8* and *35S::COP1–GFP* transgenic plants have been described previously^8,27,30,31,32^. The *hos1-3 hy5-221* double mutant was generated by crossing *hos1-3* and *hy5-221* mutants, and the *hos1-3 uvr8-6* double mutant was generated by crossing *hos1-3* and *uvr8-6* mutants. To generate the *pHY5::HY5– MYC* transgenic plants, 1.7-kb genomic fragment of *HY5* gene including a promoter sequence upstream of the translational start site was cloned into the pBA002 binary vector containing the 6× MYC sequence and the *pHY5::HY5–MYC* construct was transformed into Col-0 plants using the floral dipping method.

### UV-B tolerance assay

Seven-day-old *Arabidopsis* seedlings grown on MS-agar plates at 22□ under LDs were exposed to UV-B for appropriate durations. UV-B treatments were performed by using established conditions with broadband (Sankyo G15T8E) or narrowband UV-B lamps (Philips PL-L 36W/01/4P). The UV-B-treated seedlings were allowed to recover at 22□ for 5 d under constant light conditions. To assess the survival rate of seedlings under UV-B stress, 40∼50 seedlings were subjected to measurement survival rate. Survival was defined as plants remaining alive with green, non-wilted leaves at the end of the treatment.

### Gene expression analysis

Total RNA samples were extracted from seven-day-old seedlings grown on MS-agar plates and pretreated extensively with RNase-free DNase to get rid of genomic DNA contamination before use. cDNA was synthesized from 2 µg of total RNA using RevertAid First Strand cDNA Synthesis kit (ThermoFisher Scientific) according to the manufacturer’s recommendations. cDNAs were diluted to 60 μL with distilled water and 1 μL of diluted cDNA was used for PCR amplification.

Quantitative PCR reactions (qPCR) were performed in 96-well blocks of a StepOnePlus Real-Time PCR System (Applied Biosystems) using TOPreal SYBR Green qPCR PreMIX with low ROX (Enzynomic) in a final volume of 20 µL. Gene expression levels were normalized relative to the EUKARYOTIC TRANSLATION INITIATION FACTOR 4A1 (eIF4A) gene (At3g13920). All qPCR reactions were conducted in biological triplicates. The comparative ΔΔC_T_ approach was employed to evaluate relative quantities of individual amplified products in the samples. The threshold cycle (C_T_) for each reaction was automatically determined for each reaction by the system’s default parameters. The specificity of the RT-qPCR reactions was determined by melting curve analysis of the amplified products using the standard method installed in the system. Primers used for RT-qPCR are listed in Supplementary Table 1.

### Protein stability

Seven-day-old seedlings grown on MS-agar plates at 22□ under LDs were exposed to UV-B for varying durations. Whole seedlings were harvested for the extraction of total proteins. The MYC–HOS1 and HY5–MYC fusion proteins were immunologically detected using an anti-MYC antibody (Millipore). The HY5 proteins were immunodetected using an anti-HY5 antibody (AbClon). TUBULIN (TUB) proteins were similarly immunodetected using an anti-TUB antibody (Invitrogen) for protein quality control. The horseradish peroxidase (HRP)-conjugated secondary antibodies, goat anti-mouse IgG-HRP and goat anti-rabbit IgG-HRP (Millipore) were used for chemiluminescent western blot detection. Gel images were collected using the Fusion Molecular Imaging V15.18 software (Vilber Lourmat) and ImageQuant LAS 500 (GE Healthcare).

### RNA sequencing

Seven-day-old seedlings grown on MS-agar plates at 22□ under LDs were exposed to UV-B for 5 hours. Total RNA samples were extracted from whole seedlings using a XENOPURE PF-plant RNA purification kit (Xenohelix) according to the manufacturer’s instructions. The RNA samples were then subjected to RNA sequencing in Macrogen (Seoul, Korea). Biological triplicates were statistically analyzed. Gene ontology (GO) analysis was performed using the Biological Networks Gene Ontology tool (BiNGO) with Benjamini-Hochberg corrected *P* < 0.01. The network diagram shows significantly overrepresented GO terms. The raw RNA-seq data were deposited in the NCBI’s SRA database with accession number PRJNA1145122.

### Yeast two hybrid

To examine the physical interactions between HOS1 and other proteins, we employed yeast two-hybrid screening using the Matchmaker Gold Yeast Two-Hybrid system (Takara). The pGADT7 vector was used for GAL4 activation domain, and the pGBKT7 vector was used for GAL4 DNA-binding domain. The HOS1-coding sequences were subcloned into the pGADT7 vector and pGBKT7 vector. The UVR8, COP1, and HY5-coding sequence was subcloned into the pGADT7 vector or pGBKT7 vector. The resulting plasmids were transformed into the Y2HGold strain according to the manufacturer’s instructions. Colonies obtained from a selective medium without Leu and Trp were restreaked on a selective medium lacking Leu, Trp, Ade, and His. To prevent nonspecific growth of yeast cells, 3-amino-1,2,4-triazole was included in the media at a final concentration of 15 mM.

### Generation of polyclonal anti-HOS1 antibody in rabbit

The amplified HOS1 coding sequence was subcloned into the pCR™8/GW/TOPO vector and subsequently transferred to the gRSETA vector using the Gateway system (Invitrogen) according to the manufacturer’s instructions. The plasmid was transformed into *E. coli* BL21 (DE3). Transformed cells were grown at 37□ to an OD600 of 0.8, then induced with 1□mM isopropyl-β-D-1-thiogalactopyranoside (IPTG) for 6□h at 30□. Cells were harvested by centrifugation (6,000□rpm, 10□min), resuspended in 1x PBS, and stored at −20□. Frozen cells were thawed and subsequently incubated with 1% (v/v) Triton X-100 for 20□min, then disrupted by sonication. Soluble and insoluble fractions of His–HOS1 were separated by centrifugation at 12,000□rpm for 10□min at 4□ and analyzed by 8% SDS-PAGE. Bands corresponding to His–HOS1 in the insoluble fraction were excised from the acrylamide gel, and proteins were collected using ElectroEluter (Bio-Rad). Proteins (500□μg) were mixed with Complete Freund’s Adjuvant at a 1:1 (v/v) ratio. The prepared antigen was injected into the rabbits in three immunizations. Blood samples were collected from the immunized rabbits and centrifuged at 1,000 x □g for 10□min at 4□, and the antiserum was then purified by antigen-specific affinity chromatography using recombinant His–HOS1. His–HOS1 proteins separated by 8% SDS-PAGE were transferred to a PVDF membrane, and only the regions corresponding to His–HOS1 were excised. Membrane was blocked with 1% BSA in 1x TBS and incubated overnight at 4□ with antiserum (1:10 dilution). After three washes with 1× TBS, the membrane was sliced into 1□mm-wide blot strips, and the bound antibody was eluted with 900□μL of 0.1□M glycine (pH□2.5), followed by immediate neutralization with 100□μL of 2□M Tris–HCl (pH□8.0).

### Coimmunoprecipitation assay

Coimmunoprecipitation assays were performed as described previously^27^. Plants were grown for 7 days at 22□ and UV-B treatment. Whole plants were harvested for the assays. Plant materials were ground in liquid nitrogen, and proteins were extracted in coimmunoprecipitation buffer (50□mM Tris-Cl pH 7.4, 500□mM NaCl, 10% glycerol, 5□mM EDTA, 1% Triton-X-100, 1% Nonidet P-40, and protease inhibitor cocktail tablets (Sigma-Aldrich). Extract (5%) was used as the input control. Five μg of anti-GFP antibody (Takara) or anti-MYC (Millipore) was added to the extract and incubated for 2□h. After the incubation, protein G agarose beads were added and further incubated for 2□h. The beads were then washed five times with coimmunoprecipitation buffer lacking protease inhibitor cocktail. To elute proteins, 50□μl of 2 × SDS loading buffer (100□mM Tris-Cl pH 6.8, 4% sodium dodecyl sulfate, 0.2% bromophenol blue, 20% glycerol, 200□mM DTT) was added to the beads and incubated at 100□ for 10□min. Eluted proteins (20%) were used for IP control. Anti-GFP antibody (Takara), anti-MYC (Millipore), anti-HY5 (Abclon), anti-COP1 (Abiocode), and anti-HOS1 antibodies were used for the detection of each protein.

### Cell fractionation

Cell fraction assay was performed as described previously^28^. Seven-day-old seedlings were collected for total protein isolation in extraction buffer [20 mM Tris·HCl, pH 7.4, 25% (vol/vol) glycerol,20 mM KCl, 2 mM EDTA, 2.5 mM MgCl_2_, 250 mM sucrose, 1 mM DTT, and1 mM PMSF] at 4□. Total protein extracts were filtered through three layers of Miracloth. After centrifugation at 1,500 × g for 10 min at 4□, the clear supernatant was taken as the cytosolic fraction. The pellet was washed twice with nuclei resuspension Triton buffer [20 mM Tris·HCl, pH 7.4, 25% (vol/vol) glycerol, 2.5 mM MgCl_2_, 0.2% Triton X-100] and once with nuclei resuspension buffer [20 mM Tris·HCl, pH 7.4, 25% (vol/vol) glycerol, 2.5 mM MgCl _2_]. For protein gel blots, 30 μg of the cytosolic and 10 μg of the nuclear fraction were separated by SDS/PAGE and transferred to polyvinylidene difluoride (PVDF) membranes. Anti-UVR8 (Creative Biolabs), anti-H3 (Abcam), and anti-UGPase (Agrisera) antibodies were used to detect each protein.

### Subcellular localization analysis

Transgenic seedlings were cultivated on half-strength MS plates with 1% sucrose for 4 days. Seedlings were grown for 3 days under low-intensity white light, followed by 24 hours with (UV-B) or without (Mock) additional narrowband UV-B exposure. To stain the nuclei, individual seedlings were placed in PBS containing 5 μg/mL of 4’-6-diamidino-2-phenylindole (DAPI) (Sigma-Aldrich), covered with a coverslip, and incubated for 15 minutes before imaging. YFP–UVR8 and DAPI imaging was carried out using a Nikon AX confocal laser-scanning microscope (Nikon). DAPI and YFP were excited at wavelengths of 405 nm and 514 nm, respectively, with emissions detected between 436-482 nm using a (DAPI) and between 525-561 nm (YFP).

### Measurement of anthocyanin contents

To measure anthocyanin content, UV-B-treated seedlings were allowed to recover for 5 days at 22□ under constant light conditions. Anthocyanin was extracted by incubating the seedlings in 100% methanol at 4□ for 2 hours in complete darkness. Anthocyanin levels were analyzed using a Mithras LB940 multimode microplate reader (Berthold Technologies), with absorbance data collected through the MikroWin 2010 software (Berthold Technologies). Anthocyanin content was calculated based on absorbance values at 537, 647, and 663 nm^33^. Total anthocyanin content was determined using the equation (0.08173 × A537 - 0.00697 × A647 - 0.002228 × A663).

### Reporting Summary

Further information on research design is available in the Nature Research Reporting Summary linked to this article.

### Data availability

RNA-seq data that support the findings of this study have been deposited in National Center for Biotechnology Information Sequence Read Archive (http://www.ncbi.nlm.nih.gov/sra) under the BioProject accession number PRJNA1145122. For functional annotation of *Arabidopsis* transcripts, annotations from TAIR 10 were used (https://www.arabidopsis.org/).

