## Supplementary information for "A nuclear pore complex component HOS1 mediates UV-B light-induced UVR8 nuclear localization for activating HY5 cascade in *Arabidopsis*"

Han et al.

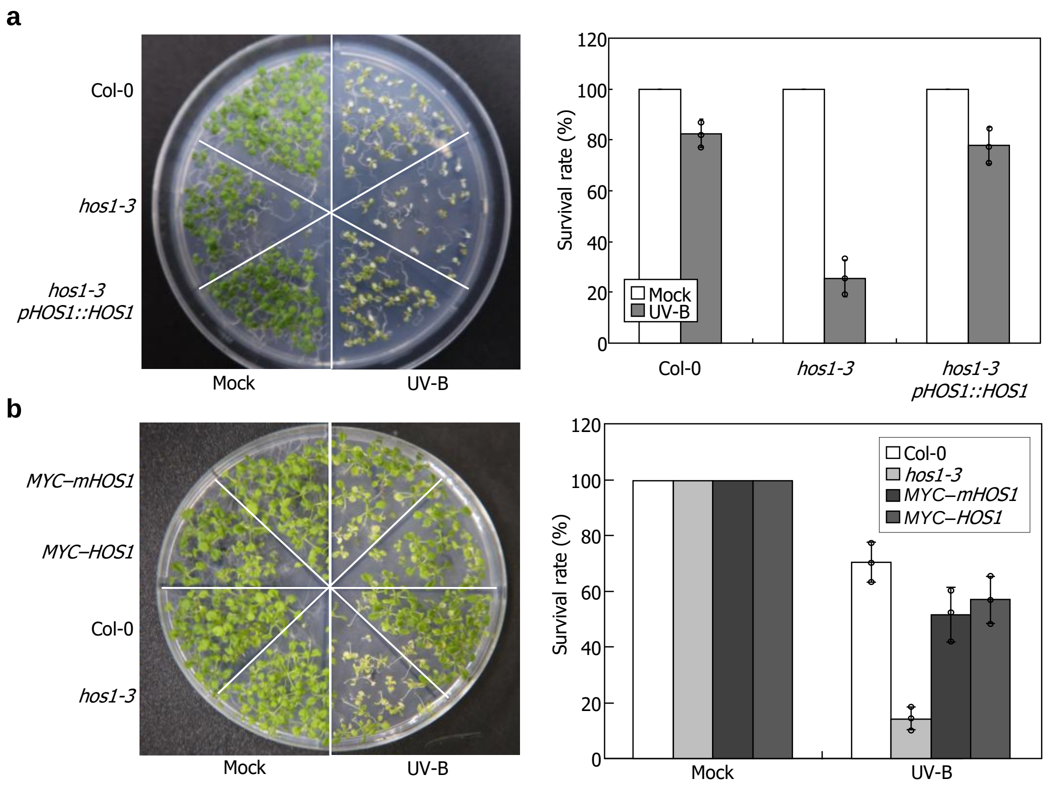


**Supplementary Fig. 1 UV-B sensitivity phenotype of *hos1-3* mutants and *HOS1* complementation lines.** Seven-day-old seedlings grown on MS-agar plates at 22℃ were exposed to UV-B radiation for 5 hours, followed by recovery for 5 days under constant light conditions at 22℃ before measuring their survival rate. Three independent measurements, each consisting of 15-20 seedlings, were statistically analyzed. Error bars indicate the standard error of the mean (SE). (**a**) A genomic complementation line (*hos1-3 pHOS1::HOS1*) expressing *HOS1* gene under the control of its native promoter in the *hos1-3* mutant background was used. (**b**) Two independent *HOS1*-overexpressing lines in the *hos1-3* mutant background expressing N-terminal MYC-tagged wild-type HOS1 protein or a modified version of HOS1 protein (mHOS1) carrying H75Y and C89S substitutions in the RING-finger domain (*MYC–HOS1* and *MYC–mHOS1*, respectively) were used.

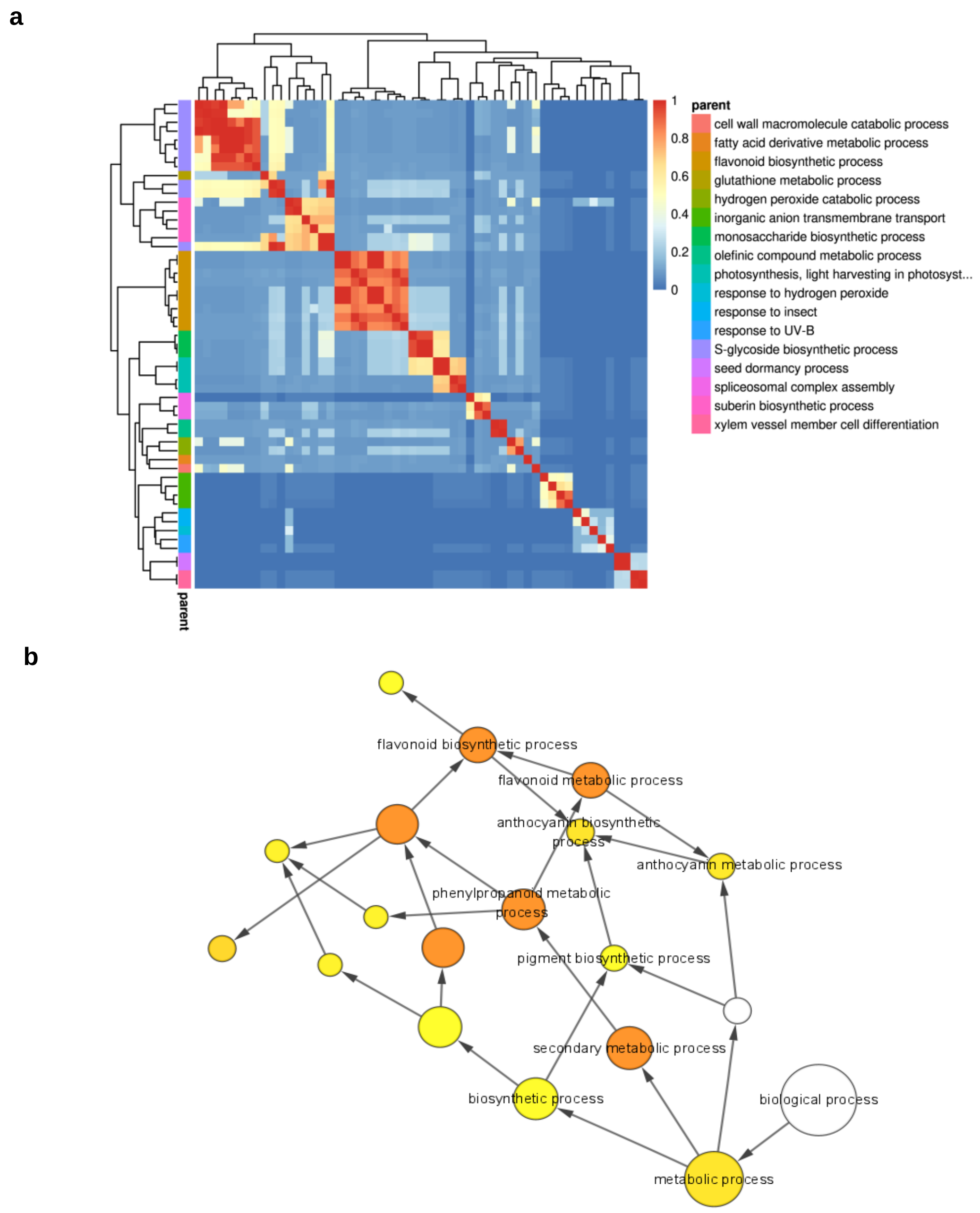


**Supplementary Fig. 2 Similarity matrix between *hos1-3* and *hy5-221* mutants after UV-B treatment.** Seven-day-old seedlings grown on MS-agar plates at 22℃ under LDs were exposed to UV-B for 5 hours and total RNA extraction from whole seedling were used in RNA sequencing analysis. Analysis criteria and conditions were described in Fig 1. The similarity matrix (**a**) and GO term analysis (**b**) illustrate that *hos1-3* and *hy5-221* mutants exhibit analogous transcriptional patterns, particularly in relation to anthocyanin synthesis and response to UV-B conditions.

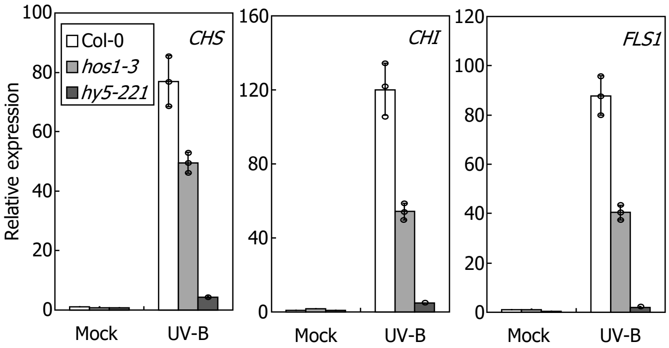


**Supplementary Fig. 3 Relative mRNA expression levels of anthocyanin biosynthesis-related genes in *hos1-3* and *hy5-221* mutants under UV-B conditions.** Plants were grown on MS-agar plates for 7 days at 22°C and exposed to UV-B for 5 hours before harvesting whole plant materials for total RNA extraction. Levels of mRNA were analyzed by RT-qPCR. Biological triplicates were statistically analyzed for each sample. Error bars indicate the standard error of the mean (SE).

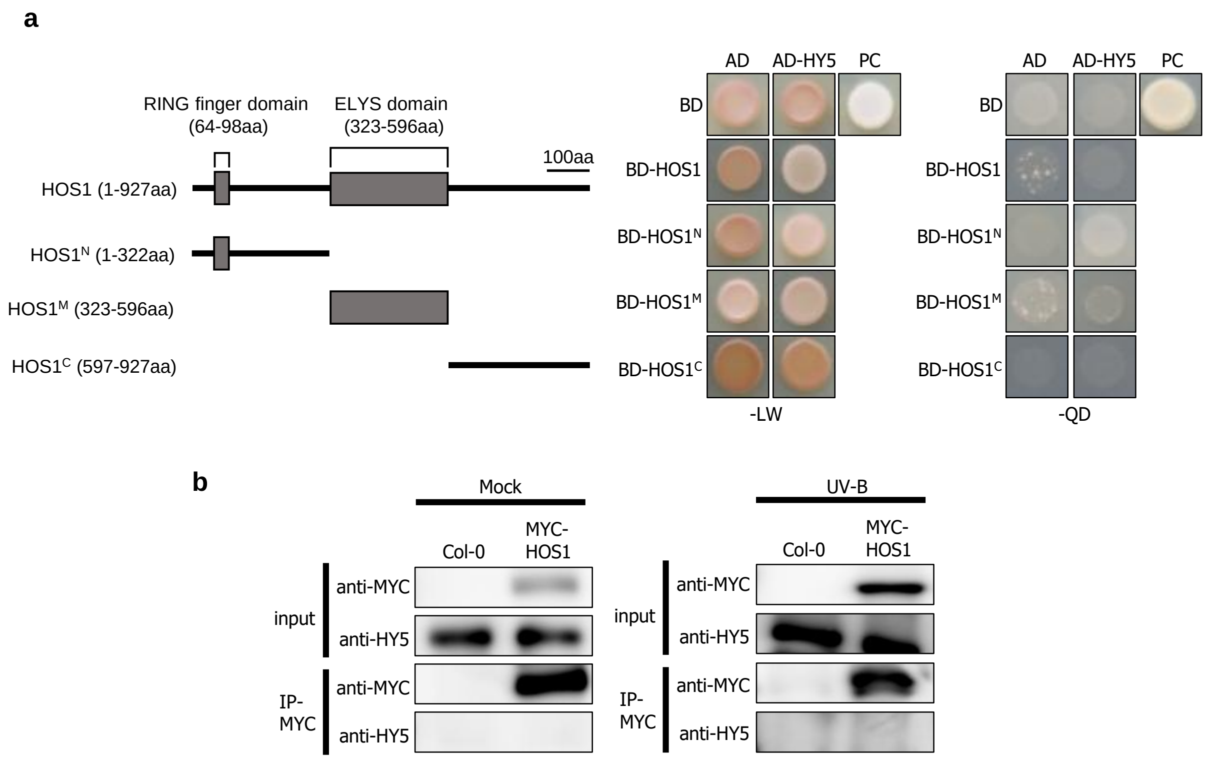


**Supplementary Fig. 4 Direct interaction between HOS1 and HY5 was not observed.** Potential interactions between HOS1 and HY5 were examined by yeast two-hybrid (Y2H) and co-immunoprecipitation (co-IP) assays. **a,** Y2H assays on HOS1–HY5 interactions in yeast cells. -LW denotes Leu and Trp dropout plates, and -QD denotes Leu, Trp, His, and Ade dropout plates. AD and BD indicates activation and DNA-binding domains, respectively. HOS1, $HOS1^N$, $HOS1^M$, and $HOS1^C$ constructs include residues 1-927, 1-323, 323-596 and 596-927, respectively. **b**, Co-IP on HOS1–HY5 interactions *in planta*. The *MYC–HOS1* transgenic plants were used for co-IP assays. Seedling growth, immuno-detection were performed, as described above Fig 3.

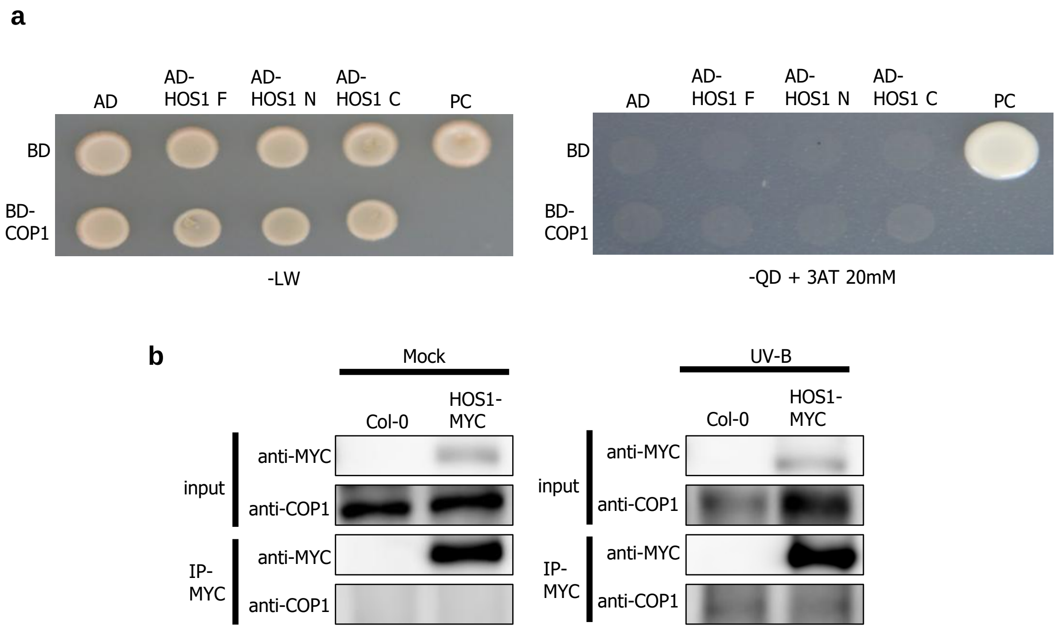


**Supplementary Fig. 5 Direct interaction between HOS1 and COP1 was not observed.** The potential interactions between HOS1 and COP1 were investigated using Y2H (**a**) and CoIP (**b**) assays. The *MYC–HOS1* transgenic plants were used for co-IP assays. TheY2H and co-IP assay were performed,  as described above.

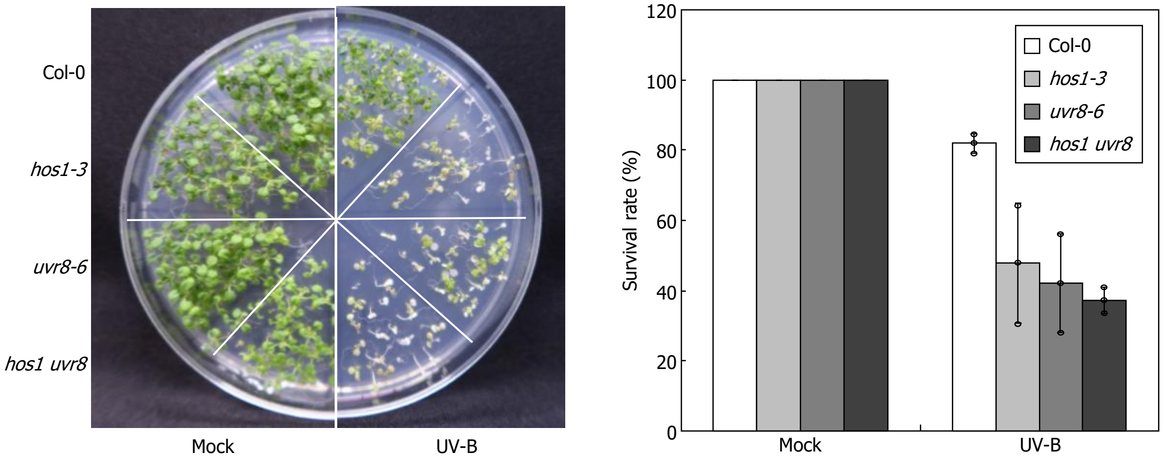


**Supplementary Fig. 6 UV-B tolerance phenotype of *hos1 uvr8* double mutants.** Seven-day-old seedlings grown on MS-agar plates at 22℃ under LDs were exposed to UV-B radiation for 5 hours. Following UV-B treatment, seedlings were allowed to recover at 22℃ under continuous light for 5 days before their survival rate was assessed. Three independent measurements, each consisting of 15-20 seedlings, were statistically analyzed. Error bars indicate the standard error of the mean (SE).

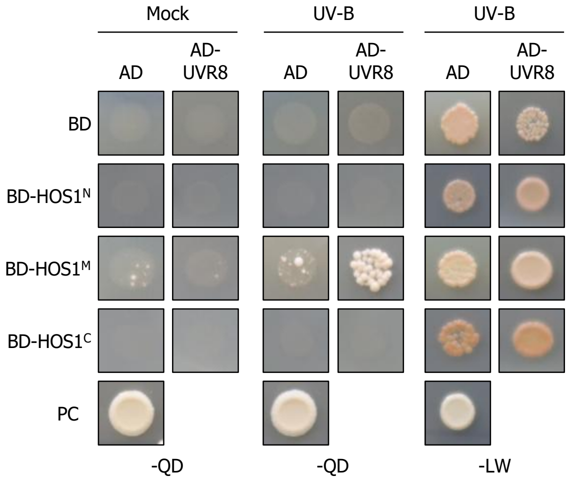


**Supplementary Fig. 7 UV-B-induced interaction between HOS1 and UVR8.** The protein–protein interactions were examined by Y2H assays. For UV-B treatment, plates were exposed to narrowband UV-B for 2 hours per day over a period of 5 days. The treatment was conducted at 30°C under dark conditions. -LW denotes Leu and Trp dropout plates, and -QD denotes Leu, Trp, His, and Ade dropout plates. AD and BD indicates activation and DNA-binding domains, respectively.

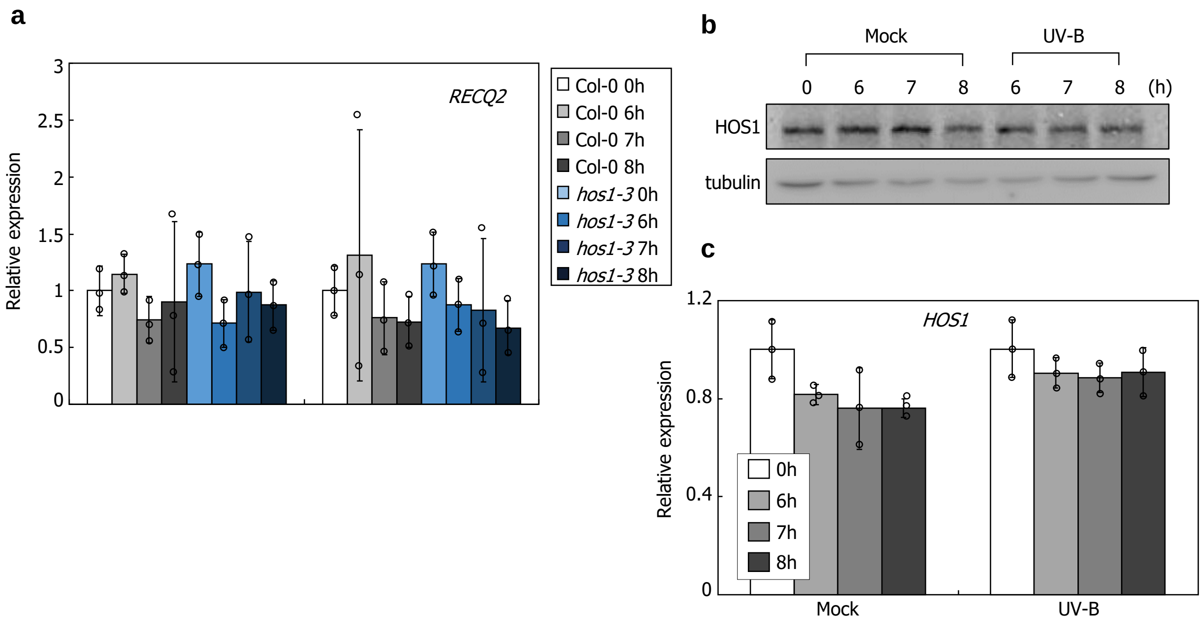


**Supplementary Fig. 8 Changes of mRNA levels of *RECQ2* and *HOS1* genes and protein levels of HOS1 after UV-B treatment.** Seven-day-old seedlings grown on MS-agar plates at 22°C were exposed to UV-B for indicated time duration. **a**, Levels of *RECQ2* mRNA in UV-B-treated seedlings. mRNA levels were analyzed by RT–qPCR. **b**, HOS1 protein levels in UV-B-treated seedlings. The *MYC–HOS1* transgenic plants were used for assays. The MYC–HOS1 proteins were immunologically detected using an anti-MYC antibody. TUB proteins were similarly detected using anti-TUB antibody for protein quality control. **c**, qRT–PCR analysis of *HOS1* mRNA levels under UV-B conditions. In **a** and **c**, biological triplicates, each consisting of 15 seedlings, were statistically analyzed.

| Primers | Sequences | Usage |
|---|---|---|
| eIF4A-F | 5’-TGACCACACAGTCTCTGCAA | RT-qPCR |
| eIF4A-R | 5’-ACCAGGGAGACTTGTTGGAC | RT-qPCR |
| CHS-F | 5’-CCAAGCTTCTTGGTCTCCGT | RT-qPCR |
| CHS-R | 5’-ACGTGCTCCACGATTGTTCT | RT-qPCR |
| CHI-F | 5’-CATCCGGAGAGTACTGCGAC | RT-qPCR |
| CHI-R | 5’-AGAGAGCTGGATTGCACCAC | RT-qPCR |
| FLS1-F | 5’-GACGGAGCTGATACGACGTT | RT-qPCR |
| FLS1-R | 5’-CCGGTTTAGCGACGGATTCT | RT-qPCR |
| HY5-F | 5’-TCGGAAAAGAAACTTCCGGT | RT-qPCR |
| HY5-R | 5’-TCCCTCGCTTCCTTTGACTT | RT-qPCR |
| RECQ2-F | 5’-GTCGAAGCGCATTTTTCCGT | RT-qPCR |
| RECQ2-R | 5’-GCTTGCGTTTCTTGCACCAT | RT-qPCR |
| HOS1-F | 5’-GCACAAGGATGCAACCAGAC | RT-qPCR |
| HOS1-R | 5’-TTGTTTCATCTGACCGCCAT | RT-qPCR |
| HOS1_Full-F | 5’-TATCCCGGGGATGGATACGAGAGAAATCAACG | Cloning |
| HOS1_Full-R | 5’-ATAGGATCCTTCATCTTGCTGCGAATCTAC | Cloning |
| HOS1_N-R | 5’-ATAGGATCCAAGCACTCTGTACACTGCAATCT | Cloning |
| HOS1_M-F | 5’-TATCCCGGGTTCCCTCTTTATTGAGGATGC | Cloning |
| HOS1_M-R | 5’-ATAGGATCCCCGAATCTAATAAGCATCTGTGC | Cloning |
| HOS1_C-F | 5’-TATCCCGGGGGCAACTGATGACCCCTC | Cloning |
| HY5_Xba1-F | 5’-TTCCCGGGTATGCAGGAACAAGCGACTA | Cloning |
| HY5_Sal1-R | 5’-CAGGTCGACGTCAAAGGCTTGCATCAGC | Cloning |
| COP1_pENTR-F | 5’-TGGTACTCGCTGCGTGAAAGGGTGGGCGCG | Cloning |
| COP1_pENTR-R | 5’-TCCGTCGAAATCTCTTCCATGGTGAAGGGGGCGGCCG | Cloning |
| UVR8_pENTR-F | 5’-CTGATGTCAAGCGTGTACGAATTTGAAAGGGTGGGCGCG | Cloning |
| UVR8_pENTR-R | 5’-CAGCCATATCCTCCGCCATGGTGAAGGGGGCGGCCG | Cloning |
| HOS1-pENTR-F | 5’-CCGCCCCCTTCACCATGGATACGAGAGAAATCAACGGTTTTG | Cloning |
| HOS1-pENTR-R | 5’-CGGCGCGCCCACCCTTTCTTGCTGCGAATCTACGTCTC | Cloning |

**Supplementary Table 1 | Primers used.**

PCR primers were designed using the NCBI Primer-BLAST tool (https://www.ncbi.nlm.nih.gov/tools/primer-blast/), ensuring that the calculated melting temperatures (Tm) of each primer pair fall within the range of 50 to 65 ℃. F and R indicate the forward and reverse primers, respectively.
